# Endosymbiont diversity across native and invasive brown widow spider populations

**DOI:** 10.1101/2023.06.22.546064

**Authors:** Monica A. Mowery, Laura C. Rosenwald, Eric Chapman, Yael Lubin, Michal Segoli, Thembile Khoza, Robin Lyle, Jennifer A. White

**Affiliations:** Mitrani Department of Desert Ecology, Blaustein Institutes for Desert Research, Ben-Gurion University of the Negev, Sede Boqer Campus, Israel; Department of Entomology, University of Kentucky, Lexington, KY, USA; South African National Biodiversity Institute, Biosystematics Division, Pretoria, South Africa; Agricultural Research Council – Plant Health and Protection, Biosystematics Division, Queenswood, South Africa

## Abstract

The invasive brown widow spider, *Latrodectus geometricus* (Araneae: Theridiidae), has spread in multiple locations around the world and, along with it, brought associated organisms such as endosymbionts. We investigated endosymbiont diversity and prevalence across putative native and invasive populations of this spider, predicting lower endosymbiont diversity across the invasive range compared to the native range. First, we characterized the microbial community in the putative native (South Africa) and invasive (Israel and the United States) ranges via high throughput 16S sequencing of 103 adult females. All specimens were dominated by reads from only 1-3 amplicon sequence variants (ASV), and most individuals were infected with an apparently uniform strain of *Rhabdochlamydia*. We also found *Rhabdochlamydia* in spider eggs, indicating that it is a maternally-inherited endosymbiont. Relatively few other ASV were detected, but included two variant *Rhabdochlamydia* strains and several *Wolbachia*, *Spiroplasma* and Enterobacteriaceae strains. We then diagnostically screened 118 adult female spiders from native and invasive populations specifically for *Rhabdochlamydia* and *Wolbachia.* We found *Rhabdochlamydia* in 86% of individuals and represented in all populations, which suggests that it is a consistent and potentially important associate of *L. geometricus. Wolbachia* was found at lower overall prevalence (14%) and was represented in all countries, but not all populations. In addition, we found evidence for geographic variation in endosymbiont prevalence: spiders from Israel were more likely to carry *Rhabdochlamydia* than those from the US and South Africa, and *Wolbachia* was geographically clustered in both Israel and South Africa. Characterizing endosymbiont prevalence and diversity is a first step in understanding their function inside the host and may shed light on the process of spread and population variability in cosmopolitan invasive species.

## Introduction

When moving into new habitats, invasive species may bring along microbial associates that can influence the invasion process (Chalkowski et al., 2018; Sepúlveda et al., 2017). Some microbes are vertically inherited endosymbionts that are restricted to the invasive species, but may yet influence interactions between the invasive and native species. Maternally-inherited endosymbionts have been shown to affect ecologically-important traits important to host fitness such as dispersal (Leonardo & Mondor, 2006), fecundity (Vorburger & Gouskov, 2011), and defenses against natural enemies (Oliver & Martinez, 2014), potentially providing an advantage to the invasive species (Jaenike, 2012). Some endosymbionts affect the organism’s reproductive biology, for example, by modifying offspring sex ratio in infected populations, which can affect the speed of invasive spread (Rey et al., 2013). For example, acting as both a mutualist and reproductive manipulator, *Rickettsia* caused whiteflies to have higher fitness and a higher proportion of daughters, and quickly spread in invasive populations (Himler et al., 2011).

Assessing endosymbiont prevalence across geographically distant populations can provide a key to understanding the role of a symbiont. Widespread prevalence of a facultative endosymbiont suggests that the symbiont plays a functional role in its host, such as providing fitness benefits or manipulating reproduction (Duron et al., 2008; Łukasik et al., 2013). The latter is often manifested by sex ratio distortions, although the most common reproductive manipulation is cytoplasmic incompatibility (CI), which causes incompatibilities between infected males and uninfected females but does not alter the sex ratio of the population (Rosenwald et al., 2020). Our understanding of the dynamics and prevalence of facultative endosymbiont infection during invasive spread is limited, especially for non-insect arthropod endosymbionts.

Invasive populations are predicted to exhibit reduced endosymbiont prevalence and diversity compared to native populations. During founding events, often few individuals are initially introduced into the invasive range (Dlugosch & Parker, 2008), in which case only a subset of endosymbionts found in the native range might be introduced to the new location (Shoemaker et al., 2000). However, in most biological invasions, multiple introductions are common (Bertelsmeier & Keller, 2018), and so endosymbiont diversity might be lower initially, and then increase over time as more individuals are introduced from various localities (Desneux et al., 2018). Comparing endosymbiont diversity across invasive and native populations can provide valuable insights into the gain and loss of microbial communities during the invasion process.

The brown widow spider, *Latrodectus geometricus* (Theridiidae), is a medically important spider with neurotoxic venom. *Latrodectus geometricus* has spread recently to multiple locations around the world from the putative native range in southern Africa, most likely via cargo shipments (Garb et al., 2004). Evidence suggests that during invasion, establishment and spread, spider traits related to dispersal, fecundity, and body size shifted across populations that were established over different time periods (Mowery, Lubin, et al., 2022). In addition to these shifts in ecologically important traits, associations with other organisms, such as parasitoids (Mowery, Arabesky, et al., 2022) or endosymbionts, may have also changed during the invasion spread.

Endosymbionts of widow spiders (genus *Latrodectus*) are poorly known. A previous study on *L. geometricus* identified the endosymbiont *Rhabdochlamydia,* but only examined a few adult females in a single, inbred lab population in Florida, USA (Dunaj et al., 2020). The same study did not detect *Rhabdochlamydia* in two other *Latrodectus* species. Hence, a further study across field-collected individuals worldwide is necessary to assess the presence of *Rhabdochlamydia* more broadly across populations of *L. geometricus*. The family *Rhabdochlamydiaceae* (Phylum: Chlamydiae) is the most diverse chlamydial family. It includes important vertebrate and human pathogens and is widespread across soil and aquatic ecosystems with many yet unknown hosts (Halter et al., 2022). The genus *Rhabdochlamydia* has been found in a few distantly-related invertebrate hosts, including a cockroach (Corsaro et al., 2007), a tick (Pillonel et al., 2019), a dwarf spider (Vanthournout & Hendrickx, 2015), and a terrestrial isopod (Kostanjšek et al., 2004), although it was not found at a high prevalence within any of these species.

Also previously found in invasive populations of *L. geometricus* was *Wolbachia,* as a facultative associate in varying prevalence across populations (Arrington, 2014). *Wolbachia* infection is common in arthropods, with 40-60% of species infected (Zug & Hammerstein, 2012), as well as in other invertebrates including nematodes (Fenn et al., 2006). *Wolbachia* is known to affects the fitness and reproduction of many of its hosts, which could have implications for successful invasive establishment and spread (Kaur et al., 2021).

In this study, we compared endosymbiont presence and diversity across populations of the brown widow spider, *L. geometricus,* from the putative native range in South Africa to populations in the invasive range in the United States and Israel, using both high-throughput sequencing and diagnostic PCR screens. Our objectives were to 1) characterize the dominant endosymbionts in *L. geometricus,* 2) compare prevalence and diversity across purported native and known invasive ranges, and 3) investigate geographic patterns of endosymbiont infection within countries. We predicted that, due to founder effects, some endosymbiont would be lost and infection rates would be lower in invasive populations in the U.S. and Israel compared to putative native populations in South Africa, and that geographic patterns of endosymbiont loss would reflect the proposed routes of invasive spread of *L. geometricus* within each country.

## Methods

### Study species

*Latrodectus geometricus,* the brown widow spider, is a globally invasive species that has established populations in parts of North and South America, the Middle East, Australia, and Asia (Sadir & Marske, 2021). In the United States, *L. geometricus* was first detected in Miami, Florida in 1936 (Pearson, 1936), was confined to southern Florida until the late 1990s, and was subsequently detected in Texas and California in the 2000s (Vincent et al., 2009). In Israel, *L. geometricus* was first detected in the Tel Aviv area in 1980 (Levy & Amitai, 1983), and in the Negev region after 2000 (Mowery *et al*., 2022). Throughout the global invasive range, *L. geometricus* is found in urban and settled habitats, and builds nests on and around buildings, on fences, garden furniture, trash bins, and in playgrounds (Sadir & Marske, 2021).

### Study sites

We collected *L. geometricus* adult females from urban environments across the United States (Edisto Island, South Carolina *n* = 10; Gainesville, Florida *n* = 10; Austin, Texas *n* = 6; Los Angeles, California *n* = 7), Israel (Haifa *n* = 7, Tel Aviv *n* = 10, Be’er Sheva *n* = 10, Yeruham *n* = 8, Midreshet Ben-Gurion *n* = 10, Eilat *n* = 1), and South Africa (Modimolle *n* = 10, Pretoria *n* = 5, Johannesburg *n* = 5, Kimberley *n* = 8, Cape Town *n* = 5, Riebeeck-Kasteel *n* = 6, George *n*= 7). Spiders were deprived of food for one week before they were preserved in 100% ethanol. Starved individuals have minimal gut content and are less likely to result in false positives for endosymbionts found in the spider’s prey (White et al., 2020). To learn about the potential for vertical transmission, we also collected *L. geometricus* egg sacs from two sites in South Africa: Kimberley (*n* = 1) and Riebeeck Kasteel (*n* = 2), and sampled egg sacs produced in the laboratory from Midreshet Ben-Gurion (*n* = 3) and Tel Aviv (*n* = 2), Israel.

### Bacterial 16S sequencing

We surface-sterilized each adult female *L. geometricus* specimen (*n* = 125) with a series of bleach and ethanol rinses (Curry et al., 2015) before longitudinally dividing the abdomen in half and extracting DNA from one half using DNeasy Blood and Tissue extraction kits (Qiagen, Germantown, MD) according to manufacturer’s instructions. In addition, we extracted DNA from the legs of two specimens, as well as from the eggs of 8 *L. geometricus* egg sacs to assess endosymbiont presence outside reproductive tissues and the potential for maternal transmission, respectively. Extraction quality for each sample was verified by PCR amplification of a segment of the COI barcoding gene (White et al., 2020). If COI failed to amplify, we attempted a second extraction with the other half of the abdomen. If this extraction failed to amplify product as well, we assumed sample preservation had been poor and eliminated the specimen from the dataset entirely (7/125 specimens). To investigate which endosymbionts were present in these specimens, we profiled the microbiomes using high-throughput sequencing of the bacterial community. We amplified the V4 region of bacterial 16S rRNA for each sample with a unique combination of indexed forward and reverse primers (Kozich et al., 2013). We visualized the resulting products, and multiplexed 1 µl aliquots from successful amplifications into one of two libraries that were purified with GenCatch PCR Cleanup Kits. Samples that failed to amplify (6/118 samples) were not included in the library. Each library also included specimens from other projects that are not reported here, and received a PhiX spike to increase sequence heterogeneity among the amplified sequences. Libraries were sequenced at the University of Kentucky genomics core facility. Sequences from each run were demultiplexed, trimmed and quality filtered within BaseSpace (Illumina, basespace.illumina.com), then imported into qiime2 (v2021.11, https://qiime2.org; Caporaso et al., 2010) using a manifest. We conducted additional quality control using deblur (Bokulich et al., 2013) implemented in qiime2 using default parameters and a trim length of 251 bases. Resulting amplicon sequence variants (ASV) were taxonomically classified using a naïve Bayes classifier that was trained on the 515F/806R V4 region of the Greengenes 13_8 99% OTUs reference database (DeSantis et al., 2006). We filtered out 15 ASV that originated from other specimens in the sequencing run (e.g., obligate endosymbionts of other host taxa, see Larsson et al., 2018 for discussion of index swapping), which collectively constituted only a small minority (0.14%) of the 3.57 x 10^6^ reads associated with the *L. geometricus* samples. Following filtering, *L. geometricus* samples with less than 1000 reads were excluded from further analysis (9/112 adult samples, 6/8 egg samples). For the remaining samples, we blasted high prevalence ASV sequences (>1% of any *L. geometricus* sample) against the NCBI nt database using the megablast algorithm, to identify bacterial taxa that may not have been included in the reference database. For ASV that appeared at very high prevalence or frequency (>90% of reads for any specimen, or found in multiple specimens across multiple locations), we amplified a longer segment of 16S using universal primers (Folmer et al., 1994) from specimen(s) dominated by that taxon, to aid in taxonomic placement.

### Diagnostic PCR

We diagnostically screened all samples (all 118 adult female, 8 egg, and 2 leg samples) for the two bacterial genera previously identified from *L. geometricus*: *Wolbachia* (Class Alphaproteobacteria, Order Rickettsiales, Family Anaplasmataceae) and *Rhabdochlamydia* (Class Chlamydia, Order Chlamydiales, Family Rhabdochlamydiaceae; Arrington, 2014; Dunaj et al., 2020). For *Wolbachia*, we used previously published protocols (Baldo et al., 2006), with primers specific to the *Wolbachia* surface protein (*wsp*) gene. For *Rhabdochlamydia*, we designed new primers in Primer3 (Untergasser et al., 2012) to amplify a ∼540bp segment of 16S: Rhabdo_108F 5’-ACACTGCCCAAACTCCTACG-3’ and Rhabdo_647R 5’-TTAGCTWCGACACAGCCAGG-3’. All reactions were run in 10µl volume; the *Rhabdochlamydia* reactions included 3 μL purified water, 1μL 10X Buffer (New England Biolabs), 1.2 μL 10mM dNTPs, 1.5 μL 25mM MgCl_2_, 0.6μL each of forward and reverse primers at 5μM, and 0.1 μL of 5U New England Biolabs Taq Polymerase. PCR reactions received one cycle of 94°C for 2 min, followed by 25 cycles of 95°C for 15 s, 56°C at 15 s, 68°C for 45 s. We electrophoresed and visualized the products on 1% agarose gels stained with Gel Red (Biotium) alongside known positive and negative (reagents-only) controls. Samples with initial negative diagnoses were retested before being categorized as uninfected. For a subset of the samples with positive evidence of infection, we repeated the PCR at a 20µl volume and purified the PCR product with either GenCatch PCR Cleanup or Gel Extraction Kits (Epoch Life Sciences, Missouri City, TX) according to manufacturer’s instructions. Products were then submitted for Sanger sequencing (Eurofins, Louisville, KY). Resulting sequences were compared to the NCBI nucleotide database using the megablast algorithm, and specimens returning a 97% or higher match to the expected bacterial genus were scored as positive. For each strain of *Wolbachia*, we sequenced 5 MLST genes (*coxA*, *fbpA*, *ftsZ*, *gatB* and *hcpA*) and the *Wolbachia* surface protein (*wsp*) according to Baldo et al. (2006). We matched resulting sequences *Wolbachia* clades (supergroups) using the PubMLST *Wolbachia* database and NCBI via the BLASTn algorithm.

For *Rhabdochlamydia,* we ran phylogenetic analyses to place the *L. geometricus* strains, using a set of accessions across Chlamydiales with *Oligosphaera ethanolica* as an outgroup. For each analysis, multiple alignments were assembled using the MAFFT server (v. 7; https://mafft.cbrc.jp/alignment/server/; Katoh et al., 2019) using the Q-INS-I alignment method that takes secondary structure into account. Maximum likelihood phylogenetic analyses were conducted on 1576-character aligned datasets using Garli (v. 2.01; Zwickl, 2006). We applied the most complex model available (GTR+I+G; Rodríguez et al., 1990) as per recommendations of Huelsenbeck and Rannala (2004) for likelihood-based analyses. We conducted a 100-replicate ML search for the tree of highest log-likelihood and a 500-replicate ML bootstrap analysis (Felsenstein, 1985) with two search replicates per individual bootstrap replicate. All analyses used the default settings.

Individual specimens were scored for the presence of *Rhabdochlamydia* and *Wolbachia* based on the combination of diagnostic, high-throughput, and Sanger sequencing data. For a sample to be scored positive, a positive diagnostic PCR needed to be corroborated by either high-throughput or Sanger sequencing validation. For a sample to be scored negative, consistent negative diagnostic PCRs needed to be accompanied by positive validation of spider COI and/or other bacterial taxa.

### Statistical methods

All analyses were conducted in R version 4.0.2. To compare the prevalence of the dominant strains of *Rhabdochlamydia* and *Wolbachia* across South Africa, Israel, and the United States, we used a general linear model (“lme4” package, Bates et al., 2015) with a binomial link function, with *Rhabdochlamydia1* or *Wolbachia1* presence or absence in an individual as the response variable, and country as the predictor.

## Results

Compared to most microbiomes in arthropods, *L. geometricus* spiders have a depauperate microbial fauna. Of 103 adult female spiders that produced sufficient read depth (mean ± SE of 33844 ± 2026 sequences per sample), all were dominated by one to three bacterial strains that accounted for greater than 90% of the reads (Figure 1). In 64 samples, a single strain accounted for greater than 99% of reads. In most samples, the most prevalent bacterial ASV was *Rhabdochlamydia* (83/103 samples) although a few samples each were dominated by ASVs corresponding to *Wolbachia* (6 samples), Enterobacteriaceae (10 samples), *Providencia* (2 samples), *Wohlfahrtimonas* (1 sample) and a bacteria that could not be placed by the Greengenes reference database, but which our analyses (see below) place within the Chlamydiales (Chlamydiales1, 2 samples, Figure 1).

**Figure 1:**
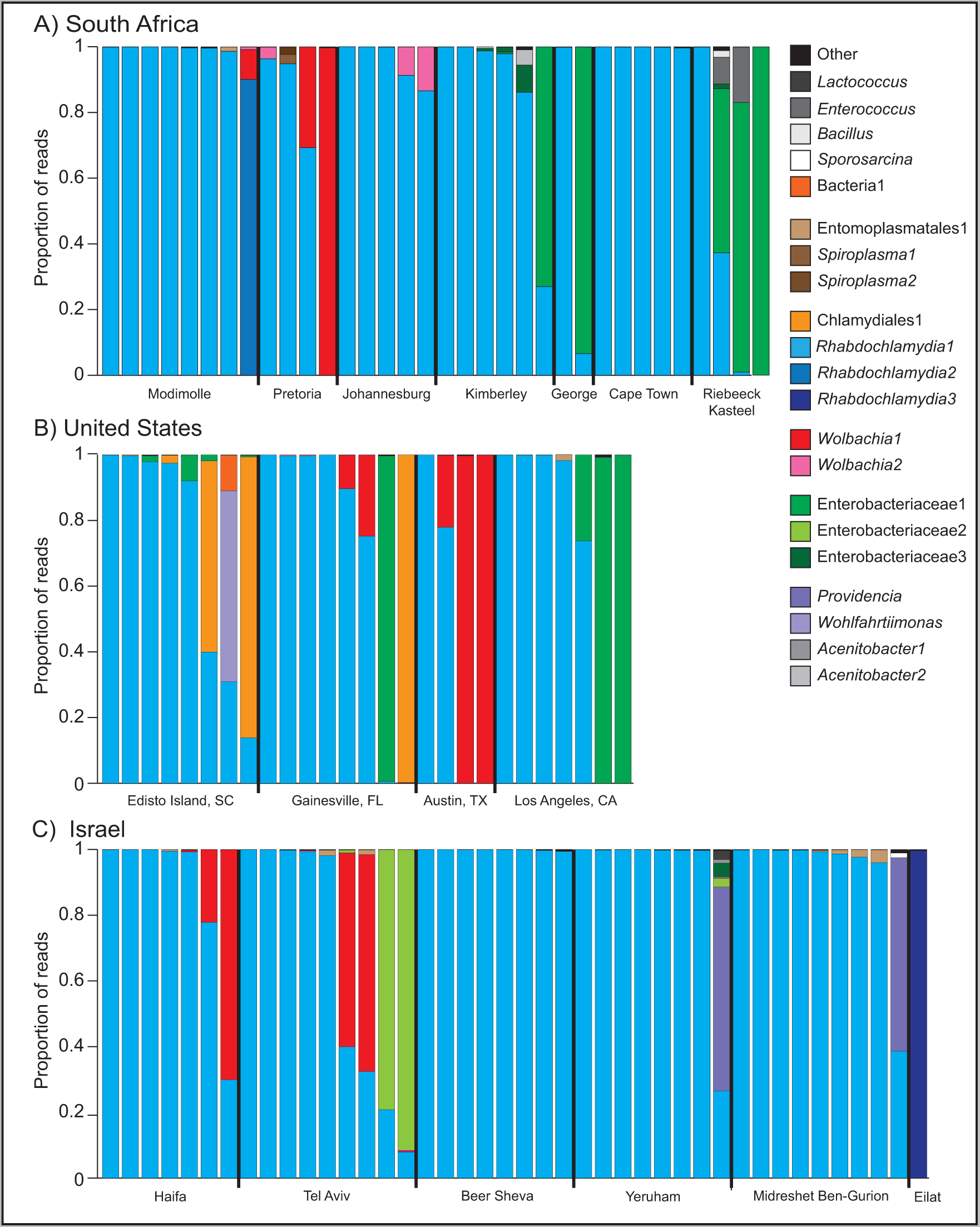
Proportional distribution of 16S sequencing reads from *Latrodectus geometricus* adult females collected from South Africa (a), the United States (b), and Israel (c). All bacterial strain types that exceeded 1% of reads in any sample are depicted. All samples that are less that 1% of reads are collected within the “other” category.

Most samples had at least some *Rhabdochlamydia* representation. Nine samples from several locations in South Africa and the United States had negligible representation (<0.1% of reads) of *Rhabdochlamydia*. The number of *Rhabdochlamydia* reads in the latter samples ranged from 0 (out of 4222 reads) to 359 (out of 37618 reads), and most fell below the number of *Rhabdochlamydia* reads seen in blanks (9-81 reads). Two samples were diagnostically positive for *Rhabdochlamydia* despite low numbers of reads, and were additionally validated by Sanger sequencing of the diagnostic product, thus were counted as *Rhabdochlamydia* positive in the final dataset. In the remaining seven specimens, the low number of proportional reads and the diagnostic absence supports the genuine absence of *Rhabdochlamydia*. Of the additional 15 samples that were excluded from high throughput analysis due to poor initial amplification or insufficient read depth, six were validated to have *Rhabdochlamydia* and nine did not.

To gain insight into the occurrence of strains of the major endosymbionts found, we used Sanger sequencing data to distinguish among strains of the same symbiont clade. Most detected *Rhabdochlamydia* strains were identical (GenBank Accession #OP598824). Two variant strains were detected, each in one individual. The variant strain from a Modimolle, South Africa specimen (#OP598825) was 99.8% similar to the dominant strain, differing at only 1/480 bases of 16S. The variant strain from Eilat, Israel (#OP598826) was 98.8% similar, differing at 6/480 bases of 16S. Phylogenetically, all three strains were clustered together within the genus *Rhabdochlamydia* and family Rhabdochlamydiaceae (Figure 2).

**Figure 2:**
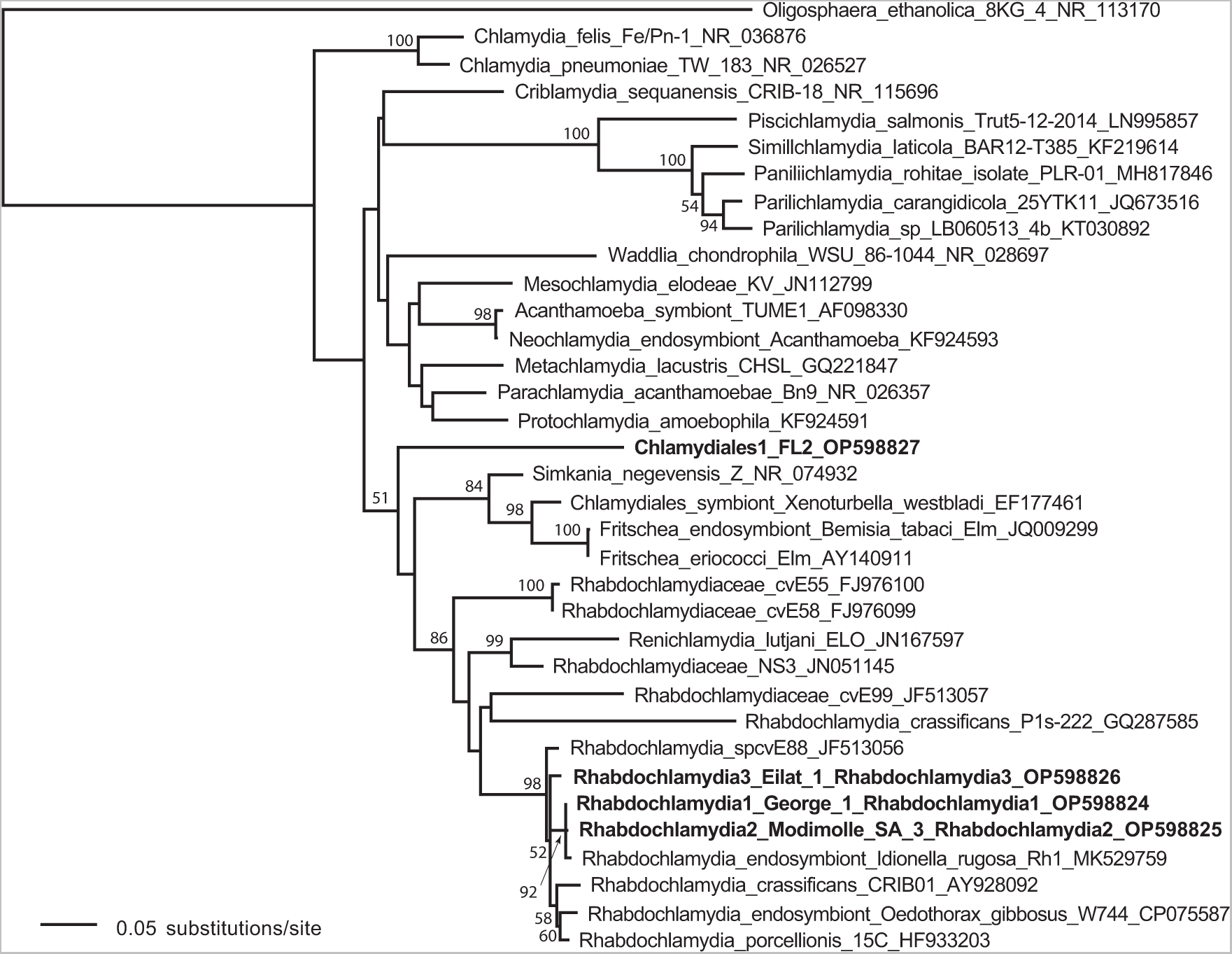
Bootstrapped phylogenetic tree of *Rhabdochlamydia* strains across spiders from the localities sampled in this study (in bold, labelled Rhabdochlamydia1, 2, and 3), Bacteria1, as well as related strains from previously published studies.

*Wolbachia* was much less common than *Rhabdochlamydia*, found in 14% (17/118) of individuals, but represented in spiders collected from all three regions. We were able to sequence all MLST genes and *wsp* for all three strains of *Wolbachia* (accession numbers OP612314-OP612330), except *gatB* in *L. geometricus Wolbachia*3. The most widespread and characteristic strain of *Wolbachia* in *L. geometricus*, *Wolbachia*1, was present in 13/118 specimens (11%), and matched most closely to Supergroup F *Wolbachia* strains (>99% similarity) based on all sequenced genes. In contrast*, L. geometricus Wolbachia*2, which was found in four specimens across three localities in South Africa, belongs to a different *Wolbachia* clade, Supergroup B, with all genes showing >99% similarity to other Supergroup B alleles. A third *Wolbachia* strain, *L. geometricus Wolbachia3,* was found in a single sample that had not been included in high throughput sequencing, but was validated with diagnostic PCR and subsequent sequencing, and appears to belong to Supergroup A. Most gene sequences were >99% similar to other supergroup A strains, although the *hcpA* gene was more divergent, bearing only 97% similarity to its closest match, a *Wolbachia* strain found in the linyphiid spider *Mermessus fradeorum*.

Only 16 other ASV, besides *Rhabdochlamydia* and *Wolbachia,* were ever found at >1% prevalence in any sample, and the majority of these (nine) were each found in single specimens. Enterobacteriaceae1 represented a substantial proportion of reads in 12 individuals across several locations in South Africa and the United States, and was the dominant ASV in eight individuals. When blasted against the NCBI database, a 1359bp segment of 16S from this bacterium (#OP598828) was not closely aligned to any other accessions, bearing greatest resemblance to aphid secondary symbionts (e.g., EU348326 at 96.8%) or *Gilliamella*, a specialized honeybee gut symbiont (e.g., CP048265 at 95.84%). Enterobacteriaceae1 was absent from Israel, although a different Enterobacteriaceae ASV was detected from two individuals collected from one location in Israel. Two other gammaproteobacteria ASVs, *Providencia* and *Wohlfahrtiimonas*, were present in two and one specimens, respectively. Bacteria1, which was found in four individuals across two locations in the southeast U.S., was not able to be placed against the Greengenes database in the qiime2 pipeline, but a 498bp segment of 16S aligns most closely with other Chlamydiales in GenBank (e.g. FJ976094 at 87.2%). Our chlamydial phylogeny (Figure 2), also supports placement within this order. Other bacterial ASV were only found at a low percentage of reads across spiders (two *Acenitobacter* ASV, two *Spiroplasma* ASV, and one each of *Entomoplasmatales, Sporosarcina, Bacillus, Enterococcus,* and *Lactococcus*.

Comparing across the three countries, a higher proportion of spiders collected in Israel were infected with the dominant strain of *Rhabdochlamydia, Rhabdochlamydia1,* than spiders from South Africa (GLM, *z* = −2.128, *p* = .033) or the U.S. (*z* = −2.538, *p* = .011). We found no differences in prevalence of the dominant *Wolbachia* strain, *Wolbachia1,* across countries (GLM, US-Israel, *z* = −0.689, *p* = .491; US-South Africa, *z* = −1.268, *p* = .205; Israel-South Africa, *z* = −0.669, *p* = .504). Using diagnostic PCR screening, we found evidence for *Rhabdochlamydia* in 100% (8/8) of *L. geometricus* eggs tested from South Africa and Israel. In contrast, only two out of eight egg sacs showed signal of *Wolbachia,* both from Tel Aviv, consistent with the proportional infection rate in adults from the source populations.

*Wolbachia* prevalence was too low for formal spatial analysis, but visually appeared to have some level of clustering (Figure 3). In South Africa, both *Wolbachia1* and *Wolbachia2* were found in northeastern populations (Johannesburg, Pretoria, and Modimolle) but were not detected elsewhere in the country. Likewise, in Israel, *Wolbachia1* was present in central and northern populations (Tel Aviv and Haifa), but was not detected in the southern Negev populations (Beer Sheva, Yeruham, Sede Boqer, Eilat). Among the four U.S. populations, *Wolbachia1* was found in spiders collected from Florida and Texas, *Wolbachia3* was in South Carolina, but no *Wolbachia* was detected in spiders from California, the most recently detected invasive population.

**Figure 3:**
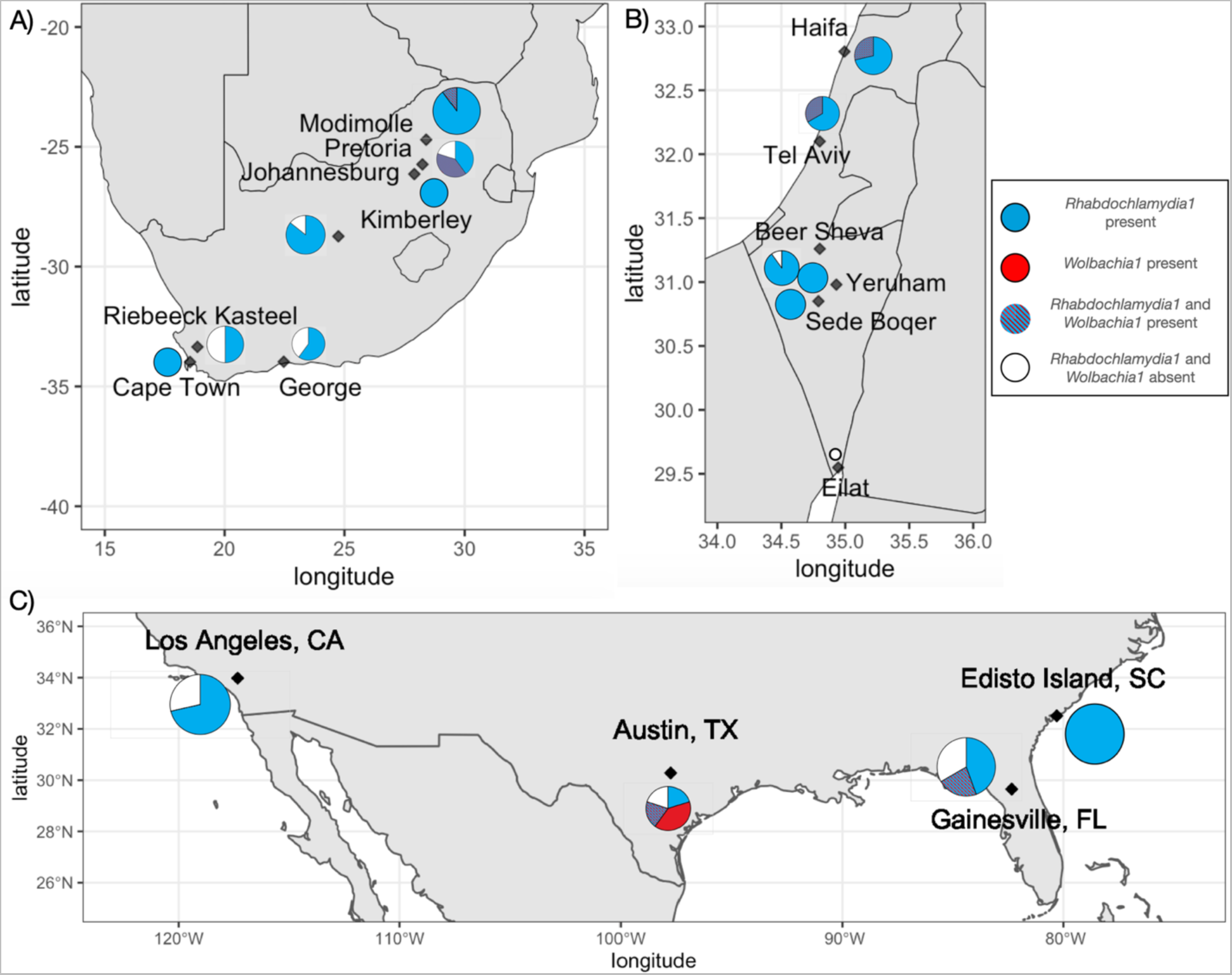
Proportion of adult female *L. geometricus* infected with *Rhabdochlamydia1* and/or *Wolbachia1* detected through PCR screening across 17 localities in A) South Africa, B) Israel, and C) the United States. Blue represents individuals infected with just *Rhabdochlamydia1,* purple represents individuals infected with both *Rhabdochlamydia1* and *Wolbachia1,* red represents individuals infected with just *Wolbachia1,* and white represents individuals infected with neither *Wolbachia1* nor *Rhabdochlamydia1.* Size of pie charts corresponds to the number of individual spiders screened from each site (range = one specimen from Eilat, Israel to 10 specimens from Edisto Island, SC, USA, see supplementary table for sample sizes and collection localities).

## Discussion

*Latrodectus geometricus* spiders have maintained a characteristic microbiome throughout their global spread. We identified one predominant endosymbiont, *Rhabdochlamydia1* in almost all spiders (86%), and represented in all collection locations. We also found a characteristic Supergroup F *Wolbachia* (*Wolbachia1*) represented in all countries, albeit in fewer individuals (11% of spiders). We detected both *Rhabdochlamydia1* and *Wolbachia1* in *L. geometricus* eggs, indicating that both are vertically transmitted endosymbionts.

The widespread presence of *Rhabdochlamydia* suggests that it might be important functionally for the host. In other arthropods, endosymbionts found at consistently high frequency across wide geographic ranges have often subsequently been found to have important fitness or reproductive consequences for their hosts (Arif et al., 2021; Cornwell & Hernández, 2021). Little is known about the functional role of *Rhabdochlamydia* in arthropods. It was described from a variety of mostly non-insect arthropods, and was generally found at low prevalence in the tested populations (Corsaro et al., 2007; Kostanjšek et al., 2004; Pillonel et al., 2019). In a terrestrial isopod, *Rhabdochlamydia* had pathogenic effects (Kostanjšek et al., 2004). The high prevalence (86%) and vertical transmission of *Rhabdochlamydia* in *L. geometricus* argue against a strongly pathogenic role for this bacterial strain within our system. *Rhabdochlamydia* was also detected in *L. geometricus* legs, which indicates that the endosymbiont is found throughout the body, and not just restricted to reproductive tissue. Genomic analysis of *Rhabdochlamydia* found in other arthropod hosts, an isopod and a tick, found pathways for polyamine synthesis (Halter et al., 2022), which are relevant for virulence and stress responses, suggesting that some strains of this bacteria are potentially beneficial in their host.

Importantly, maternal transmission also suggests the possibility of reproductive manipulation of host by symbiont. Reproductive manipulation is extremely common in vertically transmitted symbionts, and the list of bacteria that have been demonstrated to induce such manipulations is rapidly expanding (Pollmann et al., 2022; Rosenwald et al., 2020). *Rhabdochlamydia* has not yet been tested for host reproductive manipulation. The widespread prevalence and vertical transmission of *Rhabdochlamydia* in *L. geometricus* would make this system an excellent prospect for such investigations.

*Latrodectus geometricus* was also host to several strains of *Wolbachia*, a bacterial clade well known for reproductive manipulation. *Wolbachia* is common in spiders, but most strains belong to Supergroup A or B, as is the case in insects (Kaur et al., 2021). In contrast, the dominant *Wolbachia* strain in *L. geometricus* belongs to Supergroup F, which has not previously been reported for spiders. Supergroup F has been found sporadically in arthropods, including South African scorpions (Baldo et al., 2007), termites (Lo & Evans, 2007), quill mites (Glowska et al., 2015), and nematodes (Casiraghi et al., 2005). Previous work in *L. geometricus* found that *Wolbachia* likely induces mild CI, although the strain of *Wolbachia* was not characterized (Knight, 2018).

Although symbiont communities were largely similar across our sampled regions, we did find some subtle differences between the likely native and invasive ranges. *Rhabdochlamydia* was found at highest prevalence in Israel compared to populations in the U.S. and South Africa. Multiple strains of *Rhabdochlamydia, Wolbachia,* and the Enterobacteriaceae were found in South Africa, the putative native population. The dominant strain of Enterobacteriaceae was found in South Africa and the U.S., but absent in Israel, the newest invasive region that we sampled. From a previous study*, Wolbachia* prevalence in *L. geometricus* in the U.S. was highest near the initial site of introduction in Florida (Arrington, 2014). In comparison, we found lower *Wolbachia* prevalence in other locations in the southeastern and central U.S, and absence in spiders from California, the most recently established population. Similarly, in Israel, *Wolbachia* was absent in recently established populations in southern Israel. These patterns are consistent with the loss of endosymbionts during the invasion process, but more localities, specimens, and more knowledge of the invasion route is needed. Climatic differences such as hotter, dryer conditions in the Negev Desert in southern Israel could also contribute to reduction of *Wolbachia* (Charlesworth et al., 2019), although deeper sampling effort would be needed to assess whether *Wolbachia* is entirely absent from these locations.

Further work will test the functional role and fitness effects of endosymbiont presence in *L. geometricus*, as well as compare patterns of host-endosymbiont diversity during invasive spread. Invasive *L. geometricus* are highly dispersive (Mowery, Lubin, et al., 2022), and are less susceptible to parasitism by parasitoids compared to native widow species in the invasive range (Mowery, Arabesky, et al., 2022). It would be valuable to test whether these advantages and others during invasion are related to interactions with endosymbionts. In particular, the dominance and high prevalence of *Rhabdochlamydia* across global populations of *L. geometricus* suggests an important role of this endosymbiont. Characterizing potentially important and widespread endosymbionts is a step towards understanding their relevance to ecological interactions and responses to rapid environmental changes.

## Acknowledgements

We thank Ofir Altstein, Ishai Hoffmann, Cayley Buckner, Madison Heisey, Catherine Scott, Nishant Singh, Alyssa Fuller, Astri Leroy, Annari van der Merwe, Fiona Hellmann, Joh Henschel, Theresa Henschel, and Colin Ralston for assistance collecting spiders. This work was supported by a Zuckerman STEM Leadership Postdoctoral Fellowship to MAM. This material is based upon work supported by the National Science Foundation under Grant No. 1953223.

## References

Arif, S., Gerth, M., Hone-Millard, W. G., Nunes, M. D. S., Dapporto, L., & Shreeve, T. G. (2021). Evidence for multiple colonisations and *Wolbachia* infections shaping the genetic structure of the widespread butterfly *Polyommatus icarus* in the British Isles. Molecular Ecology, 30(20), 5196–5213. https://doi.org/10.1111/mec.16126

Arrington, B. D. (2014). The prevalence and effect of Wolbachia infection on the brown widow spider (Latrodectus geometricus) [Masters]. Georgia Southern University.

Baldo, L., Dunning Hotopp, J. C., Jolley, K. A., Bordenstein, S. R., Biber, S. A., Choudhury, R. R., Hayashi, C., Maiden, M. C. J., Tettelin, H., & Werren, J. H. (2006). Multilocus sequence typing system for the endosymbiont *Wolbachia pipientis*. Applied and Environmental Microbiology, 72(11), 7098–7110. https://doi.org/10.1128/AEM.00731-06

Baldo, L., Prendini, L., Corthals, A., & Werren, J. H. (2007). *Wolbachia* are present in southern African scorpions and cluster with Supergroup F. Current Microbiology, 55(5), 367–373. https://doi.org/10.1007/s00284-007-9009-4

Bates, D., Mächler, M., Bolker, B., & Walker, S. (2015). Fitting Linear Mixed-Effects Models using lme4. Journal of Statistical Software, 67(1), Article 1. https://doi.org/10.18637/jss.v067.i01

Bertelsmeier, C., & Keller, L. (2018). Bridgehead effects and role of adaptive evolution in invasive populations. Trends in Ecology & Evolution, 33(7), 527–534. https://doi.org/10.1016/j.tree.2018.04.014

Bokulich, N. A., Subramanian, S., Faith, J. J., Gevers, D., Gordon, J. I., Knight, R., Mills, D. A., & Caporaso, J. G. (2013). Quality-filtering vastly improves diversity estimates from Illumina amplicon sequencing. Nature Methods, 10(1), 57–59. https://doi.org/10.1038/nmeth.2276

Caporaso, J. G., Kuczynski, J., Stombaugh, J., Bittinger, K., Bushman, F. D., Costello, E. K., Fierer, N., Peña, A. G., Goodrich, J. K., Gordon, J. I., Huttley, G. A., Kelley, S. T., Knights, D., Koenig, J. E., Ley, R. E., Lozupone, C. A., McDonald, D., Muegge, B. D., Pirrung, M., … Knight, R. (2010). QIIME allows analysis of high-throughput community sequencing data. Nature Methods, 7(5), 335–336. https://doi.org/10.1038/nmeth.f.303

Casiraghi, M., Bordenstein, S. R., Baldo, L., Lo, N., Beninati, T., Wernegreen, J. J., Werren, J. H., & Bandi, C. Y. 2005. (2005). Phylogeny of *Wolbachia pipientis* based on gltA, groEL and ftsZ gene sequences: Clustering of arthropod and nematode symbionts in the F supergroup, and evidence for further diversity in the *Wolbachia* tree. Microbiology, 151(12), 4015–4022. https://doi.org/10.1099/mic.0.28313-0

Chalkowski, K., Lepczyk, C. A., & Zohdy, S. (2018). Parasite ecology of invasive species: Conceptual framework and new hypotheses. Trends in Parasitology, 34(8), 655– 663. https://doi.org/10.1016/j.pt.2018.05.008

Charlesworth, J., Weinert, L. A., Araujo, E. V., & Welch, J. J. (2019). *Wolbachia*, *Cardinium* and climate: An analysis of global data. Biology Letters, 15(8), 20190273. https://doi.org/10.1098/rsbl.2019.0273

Cornwell, B. H., & Hernández, L. (2021). Genetic structure in the endosymbiont *Breviolum ‘muscatinei’* is correlated with geographical location, environment and host species. Proceedings of the Royal Society B: Biological Sciences, 288(1946), 20202896. https://doi.org/10.1098/rspb.2020.2896

Corsaro, D., Thomas, V., Goy, G., Venditti, D., Radek, R., & Greub, G. (2007). ‘Candidatus *Rhabdochlamydia crassificans*’, an intracellular bacterial pathogen of the cockroach *Blatta orientalis* (Insecta: Blattodea). Systematic and Applied Microbiology, 30(3), 221–228. https://doi.org/10.1016/j.syapm.2006.06.001

Curry, M. M., Paliulis, L. V., Welch, K. D., Harwood, J. D., & White, J. A. (2015). Multiple endosymbiont infections and reproductive manipulations in a linyphiid spider population. Heredity, 115(2), Article 2. https://doi.org/10.1038/hdy.2015.2

DeSantis, T. Z., Hugenholtz, P., Larsen, N., Rojas, M., Brodie, E. L., Keller, K., Huber, T., Dalevi, D., Hu, P., & Andersen, G. L. (2006). Greengenes, a chimera-checked 16S rRNA gene database and workbench compatible with ARB. Applied and Environmental Microbiology, 72(7), 5069–5072. https://doi.org/10.1128/AEM.03006-05

Desneux, N., Asplen, M. K., Brady, C. M., Heimpel, G. E., Hopper, K. R., Luo, C., Monticelli, L., Oliver, K. M., & White, J. A. (2018). Intraspecific variation in facultative symbiont infection among native and exotic pest populations: Potential implications for biological control. Biological Control, 116, 27–35. https://doi.org/10.1016/j.biocontrol.2017.06.007

Dlugosch, K. M., & Parker, I. M. (2008). Founding events in species invasions: Genetic variation, adaptive evolution, and the role of multiple introductions. Molecular Ecology, 17(1), 431–449. https://doi.org/10.1111/j.1365-294X.2007.03538.x

Dunaj, S. J., Bettencourt, B. R., Garb, J. E., & Brucker, R. M. (2020). Spider phylosymbiosis: Divergence of widow spider species and their tissues’ microbiomes. BMC Evolutionary Biology, 20(1), 104. https://doi.org/10.1186/s12862-020-01664-x

Duron, O., Hurst, G. D. D., Hornett, E. A., Josling, J. A., & Engelstädter, J. (2008). High incidence of the maternally inherited bacterium *Cardinium* in spiders. Molecular Ecology, 17(6), 1427–1437. https://doi.org/10.1111/j.1365-294X.2008.03689.x

Felsenstein, J. (1985). Confidence limits on phylogenies: An approach using the bootstrap. Evolution, 39(4), 783–791. https://doi.org/10.2307/2408678

Fenn, K., Conlon, C., Jones, M., Quail, M. A., Holroyd, N. E., Parkhill, J., & Blaxter, M. (2006). Phylogenetic relationships of the *Wolbachia* of nematodes and arthropods. PLOS Pathogens, 2(10), e94. https://doi.org/10.1371/journal.ppat.0020094

Folmer, O., Black, M., Hoeh, W., Lutz, R., & Vrijenhoek, R. (1994). DNA primers for amplification of mitochondrial cytochrome c oxidase subunit I from diverse metazoan invertebrates. Molecular Marine Biology and Biotechnology, 3(5), 294–299.

Garb, J. E., González, A., & Gillespie, R. G. (2004). The black widow spider genus *Latrodectus* (Araneae: Theridiidae): Phylogeny, biogeography, and invasion history. Molecular Phylogenetics and Evolution, 31(3), 1127–1142. https://doi.org/10.1016/j.ympev.2003.10.012

Glowska, E., Dragun-Damian, A., Dabert, M., & Gerth, M. (2015). New *Wolbachia* supergroups detected in quill mites (Acari: Syringophilidae). Infection, Genetics and Evolution, 30, 140–146. https://doi.org/10.1016/j.meegid.2014.12.019

Halter, T., Köstlbacher, S., Collingro, A., Sixt, B. S., Tönshoff, E. R., Hendrickx, F., Kostanjšek, R., & Horn, M. (2022). Ecology and evolution of chlamydial symbionts of arthropods. ISME Communications, 2(1), Article 1. https://doi.org/10.1038/s43705-022-00124-5

Hilgenboecker, K., Hammerstein, P., Schlattmann, P., Telschow, A., & Werren, J. H. (2008). How many species are infected with *Wolbachia*?—A statistical analysis of current data. FEMS Microbiology Letters, 281(2), 215–220. https://doi.org/10.1111/j.1574-6968.2008.01110.x

Himler, A. G., Adachi-Hagimori, T., Bergen, J. E., Kozuch, A., Kelly, S. E., Tabashnik, B. E., Chiel, E., Duckworth, V. E., Dennehy, T. J., Zchori-Fein, E., & Hunter, M. S. (2011). Rapid spread of a bacterial symbiont in an invasive whitefly is driven by fitness benefits and female bias. Science, 332(6026), 254–256. https://doi.org/10.1126/science.1199410

Huelsenbeck, J. P., & Rannala, B. (2004). Frequentist properties of Bayesian posterior probabilities of phylogenetic trees under simple and complex substitution models. Systematic Biology, 53(6), 904–913. https://doi.org/10.1080/10635150490522629

Jaenike, J. (2012). Population genetics of beneficial heritable symbionts. Trends in Ecology & Evolution, 27(4), 226–232. https://doi.org/10.1016/j.tree.2011.10.005

Katoh, K., Rozewicki, J., & Yamada, K. D. (2019). MAFFT online service: Multiple sequence alignment, interactive sequence choice and visualization. Briefings in Bioinformatics, 20(4), 1160–1166. https://doi.org/10.1093/bib/bbx108

Kaur, R., Shropshire, J. D., Cross, K. L., Leigh, B., Mansueto, A. J., Stewart, V., Bordenstein, S. R., & Bordenstein, S. R. (2021). Living in the endosymbiotic world of *Wolbachia*: A centennial review. Cell Host & Microbe, 29(6), 879–893. https://doi.org/10.1016/j.chom.2021.03.006

Knight, E. (2018). Characterizing the complex relationship between the brown widow spider and Its bacterial endosymbiont, *Wolbachia*. Electronic Theses and Dissertations. https://digitalcommons.georgiasouthern.edu/etd/1825

Kostanjšek, R., Štrus, J., Drobne, D., & Avguštin, G. (2004). ‘Candidatus *Rhabdochlamydia porcellionis*’, an intracellular bacterium from the hepatopancreas of the terrestrial isopod *Porcellio scaber* (Crustacea: Isopoda). International Journal of Systematic and Evolutionary Microbiology, 54(2), 543–549. https://doi.org/10.1099/ijs.0.02802-0

Kozich, J. J., Westcott, S. L., Baxter, N. T., Highlander, S. K., & Schloss, P. D. (2013). Development of a dual-index sequencing strategy and curation pipeline for analyzing amplicon sequence data on the MiSeq Illumina sequencing platform. Applied and Environmental Microbiology, 79(17), 5112–5120. https://doi.org/10.1128/AEM.01043-13

Larsson, A., Stanley, G. M., Sinha, R., Weissman, I., & Sandberg, R. (2018). Computational correction of index switching in multiplexed sequencing libraries. Nature Methods. https://doi.org/10.1038/nmeth.4666

Leonardo, T. E., & Mondor, E. B. (2006). Symbiont modifies host life-history traits that affect gene flow. Proceedings of the Royal Society B: Biological Sciences, 273(1590), 1079–1084. https://doi.org/10.1098/rspb.2005.3408

Levy, G., & Amitai, P. (1983). Revision of the widow-spider genus *Latrodectus* (Araneae: Theridiidae) in Israel. Zoological Journal of the Linnean Society, 77(1), 39–63. https://doi.org/10.1111/j.1096-3642.1983.tb01720.x

Lo, N., & Evans, T. A. (2007). Phylogenetic diversity of the intracellular symbiont *Wolbachia* in termites. Molecular Phylogenetics and Evolution, 44(1), 461–466. https://doi.org/10.1016/j.ympev.2006.10.028

Łukasik, P., van Asch, M., Guo, H., Ferrari, J., & Charles J. Godfray, H. (2013). Unrelated facultative endosymbionts protect aphids against a fungal pathogen. Ecology Letters, 16(2), 214–218. https://doi.org/10.1111/ele.12031

Mowery, M. A., Arabesky, V., Lubin, Y., & Segoli, M. (2022). Differential parasitism of native and invasive widow spider egg sacs. Behavioral Ecology, 33(3), 565–572. https://doi.org/10.1093/beheco/arac017

Mowery, M. A., Lubin, Y., Harari, A., Mason, A. C., & Andrade, M. C. B. (2022). Dispersal and life history of brown widow spiders in dated invasive populations on two continents. Animal Behaviour. https://doi.org/10.1016/j.anbehav.2022.02.006

Oliver, K. M., & Martinez, A. J. (2014). How resident microbes modulate ecologically-important traits of insects. Current Opinion in Insect Science, 4, 1–7. https://doi.org/10.1016/j.cois.2014.08.001

Pearson, J. F. W. (1936). *Latrodectus geometricus* Koch in southern Florida. Science, 83(2161), 522–523.

Pillonel, T., Bertelli, C., Aeby, S., de Barsy, M., Jacquier, N., Kebbi-Beghdadi, C., Mueller, L., Vouga, M., & Greub, G. (2019). Sequencing the obligate intracellular *Rhabdochlamydia helvetica* within its tick host *Ixodes ricinus* to investigate their symbiotic relationship. Genome Biology and Evolution, 11(4), 1334–1344. https://doi.org/10.1093/gbe/evz072

Pollmann, M., Moore, L. D., Krimmer, E., D’Alvise, P., Hasselmann, M., Perlman, S. J., Ballinger, M. J., Steidle, J. L. M., & Gottlieb, Y. (2022). Highly transmissible cytoplasmic incompatibility by the extracellular insect symbiont *Spiroplasma*. IScience, 25(5), 104335. https://doi.org/10.1016/j.isci.2022.104335

Rey, O., Estoup, A., Facon, B., Loiseau, A., Aebi, A., Duron, O., Vavre, F., & Foucaud, J. (2013). Distribution of endosymbiotic reproductive manipulators reflects invasion process and not reproductive system polymorphism in the little fire ant *Wasmannia auropunctata*. PLOS ONE, 8(3), e58467. https://doi.org/10.1371/journal.pone.0058467

Rodríguez, F., Oliver, J. L., Marín, A., & Medina, J. R. (1990). The general stochastic model of nucleotide substitution. Journal of Theoretical Biology, 142(4), 485–501. https://doi.org/10.1016/S0022-5193(05)80104-3

Rosenwald, L. C., Sitvarin, M. I., & White, J. A. (2020). Endosymbiotic *Rickettsiella* causes cytoplasmic incompatibility in a spider host. *Proceedings*. Biological Sciences, 287(1930), 20201107. https://doi.org/10.1098/rspb.2020.1107

Sadir, M., & Marske, K. A. (2021). Urban environments aid invasion of brown widows (Theridiidae: *Latrodectus geometricus*) in North America, constraining regions of overlap and mitigating potential impact on native widows. Frontiers in Ecology and Evolution, 9. https://www.frontiersin.org/articles/10.3389/fevo.2021.757902

Sepúlveda, D. A., Zepeda-Paulo, F., Ramírez, C. C., Lavandero, B., & Figueroa, C. C. (2017). Diversity, frequency, and geographic distribution of facultative bacterial endosymbionts in introduced aphid pests. Insect Science, 24(3), 511–521. https://doi.org/10.1111/1744-7917.12313

Shoemaker, D. D., Ross, K. G., Keller, L., Vargo, E. L., & Werren, J. H. (2000). *Wolbachia* infections in native and introduced populations of fire ants (*Solenopsis spp*.). Insect Molecular Biology, 9(6), 661–673. https://doi.org/10.1046/j.1365-2583.2000.00233.x

Untergasser, A., Cutcutache, I., Koressaar, T., Ye, J., Faircloth, B. C., Remm, M., & Rozen, S. G. (2012). Primer3—New capabilities and interfaces. Nucleic Acids Research, 40(15), e115. https://doi.org/10.1093/nar/gks596

Vanthournout, B., & Hendrickx, F. (2015). Endosymbiont dominated bacterial communities in a dwarf spider. PLOS ONE, 10(2), e0117297. https://doi.org/10.1371/journal.pone.0117297

Vincent, L. S., Vetter, R. S., Wrenn, W. J., Kempf, J. K., & Berrian, J. E. (2009). The brown widow spider *Latrodectus geometricus* C. L. Koch, 1841, in southern California. The Pan-Pacific Entomologist, 84(4), 344–349. https://doi.org/10.3956/2008-07.1

Vorburger, C., & Gouskov, A. (2011). Only helpful when required: A longevity cost of harbouring defensive symbionts. Journal of Evolutionary Biology, 24(7), 1611–1617. https://doi.org/10.1111/j.1420-9101.2011.02292.x

White, J. A., Styer, A., Rosenwald, L. C., Curry, M. M., Welch, K. D., Athey, K. J., & Chapman, E. G. (2020). Endosymbiotic bacteria are prevalent and diverse in agricultural spiders. Microbial Ecology, 79(2), 472–481. https://doi.org/10.1007/s00248-019-01411-w

Xie, J., Butler, S., Sanchez, G., & Mateos, M. (2014). Male killing *Spiroplasma* protects *Drosophila melanogaster* against two parasitoid wasps. Heredity, 112(4), Article 4. https://doi.org/10.1038/hdy.2013.118

Zug, R., & Hammerstein, P. (2012). Still a host of hosts for *Wolbachia*: Analysis of recent data suggests that 40% of terrestrial arthropod species are infected. PLOS ONE, 7(6), e38544. https://doi.org/10.1371/journal.pone.0038544

Zwickl, D. J. (2006). Genetic algorithm approaches for the phylogenetic analysis of large biological sequence datasets under the maximum likelihood criterion [Ph.D., The University of Texas at Austin]. https://www.proquest.com/docview/304984400/abstract/CA8369C6EFDA4D21PQ/1

